# Hsa-mir-210 as a novel signature predicts survival in lung squamous cell carcinoma using bioinformatics analysis

**DOI:** 10.1101/660175

**Authors:** Yu Liu, Yu Chen, Ling-Ling Li, Shen-Nan Wang, Shu Xia

**Author notes:** Correspondence to: Shu Xia, Department of Oncology, Tongji Hospital, Tongji Medical College, Huazhong University of Science and Technology, 1095 Jiefang Avenue, Wuhan, P.R. China.

## Abstract

In recent years, more and more studies have shown that the expression of miRNAs is closely related to the occurrence of tumors and plays an irreplaceable role in the development and metastasis of tumors. The research was focused on lung squamous cell carcinoma. The data information is downloaded from the TCGA database and analyzed for variance, which is then verified in the GEO database. Then differential expression of miRNAs was found and survival analysis was performed, through the cut – off standard(P<0.05,|logFC|≥2), we screen the 38 up-regulated miRNAs and 14 down-regulated miRNAs from the TCGA. Finally, after the verification on the GEO database, four up-regulated miRNAs (hsa-mir-205, hsa-mir-210, hsa-mir-182, hsa-mir-224) and one down-regulated miRNA (hsa-mir-451) were obtained, which provide a new direction for the diagnosis of lung squamous cell carcinoma. In the survival analysis, it was found that the expression state of hsamir-210 was significantly correlated with patient survival. The results in the univariate and multivariate Cox analysis indicated that hsa-mir-210 was an independent prognostic factor on lung squamous cell carcinoma. The functional enrichment analysis showed that hsa-mir-210 was closely related to positive and negative regulation of cell proliferation, DNA transcription, VEGF signaling pathway, MAPK signaling pathway, hif-1 signaling pathway and choline metabolic pathway. In summary, this study suggested that hsa-mir-210 could be a potential prognostic factor for lung squamous cell carcinoma.

**Author Summary:** MicroRNAs are single-stranded small molecular RNA that participate in the regulation of various biological functions through indirect regulation of gene expression, and have been reported to play an important role in the occurrence, development, invasion and metastasis of tumors. In recent years, the research on miRNAs has become increasingly hot, and the role of miRNAs in tumor has been proved more and more. The subjects of this study were squamous cell carcinoma in lung cancer with relatively few studies on miRNAs. Through high-throughput data analysis, miRNAs with differential expression between lung squamous cell carcinoma tissues and normal tissues were found. These differentially expressed miRNAs provide a new direction for the early diagnosis of patients. Then, survival analysis was conducted to find the miRNAs significantly correlated with the total survival time of patients, and multi-factor analysis was conducted to exclude the influence of other clinical factors, and independent risk factors (miRNAs) affecting the survival of patients were determined, so as to provide new targets for the treatment and survival prediction of patients.

## Introduction

Lung cancer has the highest mortality rate of all cancers and is the leading cause of cancer-related deaths worldwide [1, 2]. Most (85%) of lung cancers are classified as non-small-cell lung cancer (NSCLC) and small-cell lung cancer (15%) (SCLC). The two predominant NSCLC histological phenotypes are adenocarcinoma and squamous cell carcinoma (LUSC)[3]. LUSC is associated with greater mortality and morbidity due to its highly invasive nature that often invades neighboring tissues, and can metastasize distant organs[4].The treatments of the LUSC are often less effective and the chemotherapy remains the major therapeutic choice[5].The mortality is particularly higher in advanced stages compared with early interventions. At the present, LUSC still lacks effective molecular target for developing target therapy. Therefore, it is necessary to explore novel biomarkers for prognosis prediction and development of molecular target therapy for LUSC patients [1, 6].

MicroRNAs (miRNAs), a key component of the small and noncoding RNA family, are approximately 18–25nucleotides that involved in the post-transcriptional regulation of gene expression[7]. By binding to the 3’ or 5’ untranslated region of the target transcripts, miRNAs can modulate genes expression through translational repression or cleavage of mRNA [7, 8]. It has been shown that miRNAs are aberrantly expressed in various types of malignancies and function either as oncogenes or tumor suppressors. Accumulating evidence has demonstrated that miRNAs regulated various carcinogenesis processes including cell maturation, cell proliferation, migration, invasion, autophagy, apoptosis, and metastasis[9]. Therefore, this suggests that miRNAs can be potential noninvasive biomarkers for cancer[8] and have a large potential to serve as promising markers in the diagnosis, prognosis, and personalized targeted therapies[9, 10].

Although there are a large number of studies on the relationship between miRNAs and lung cancer, there are also some marker molecules that were identified to predict clinical survivals. However, many studies have focused on lung cancer or NSCLS, and there are few studies on the subclass of squamous cell carcinoma. And there are many inconsistence exist in previous studies which may due to the small sample size, heterogeneous histological subtype, different detection platforms, and various data processing methods. The Cancer Genome Atlas (TCGA) is a large-scale, collaborative effort led by the National Cancer Institute and the National Human Genome Research Institute to map the genomic and epigenetic changes that occur in 32 types of human cancer, including nine rare tumors[11]. There are a large number of high-throughput miRNAs sequencing data on LUSC in TCGA. The miRNAs sequencing data of LUSC and normal tissues used in this study were downloaded from the TCGA database. The clinical data of the sample is also obtained in TCGA. By analyzing differentially expressed miRNAs and verify in the Gene Expression Omnibus (GEO) database, we find a correlation with patient survival and identify miRNA that can effectively predict patient survival. On this basis, we analyze the protein expression and biological function of potential target miRNAs, and provide a new understanding of the molecular mechanism of LUSC.

## Results

The main research data of TCGA obtained 387 samples of miRNAs expression information, including 342 LUSC tissue samples and 45 normal tissue samples (S1 Table), and obtained clinical information of 337 samples (S2 Table), mainly including diagnosis age, sex, smoking history category, metastasis, lymph node stage, tumor stage, etc. (Table 1). According to the cut-off criteria, under the condition of P<0.05, the expression was up-regulated by logFC≥2, and the expression of logFC≤-2 was down-regulated. 38 up-regulated miRNAs were expressed and 14 down-regulated miRNAs were expressed (Table2). In order to understand the expression of the miRNAs and whether the P value and the logFC are logical, we show the result as a volcano map (Fig1).

**Fig 1.**
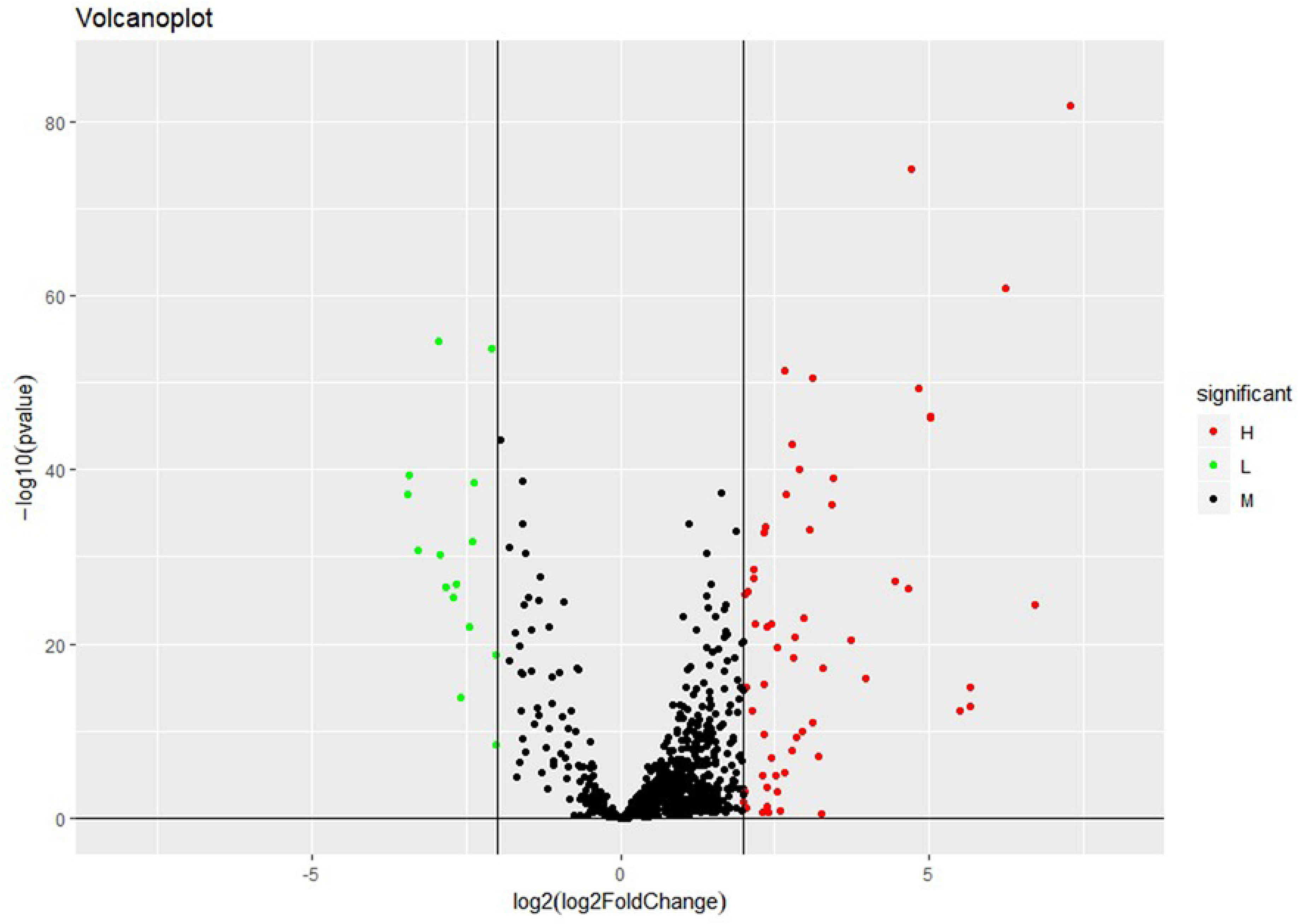
Volcano plot of differentially expressed miRNAs. The red dot represents up-regulated miRNAs, and green dot represents down-regulated miRNAs.

**Table 1.**
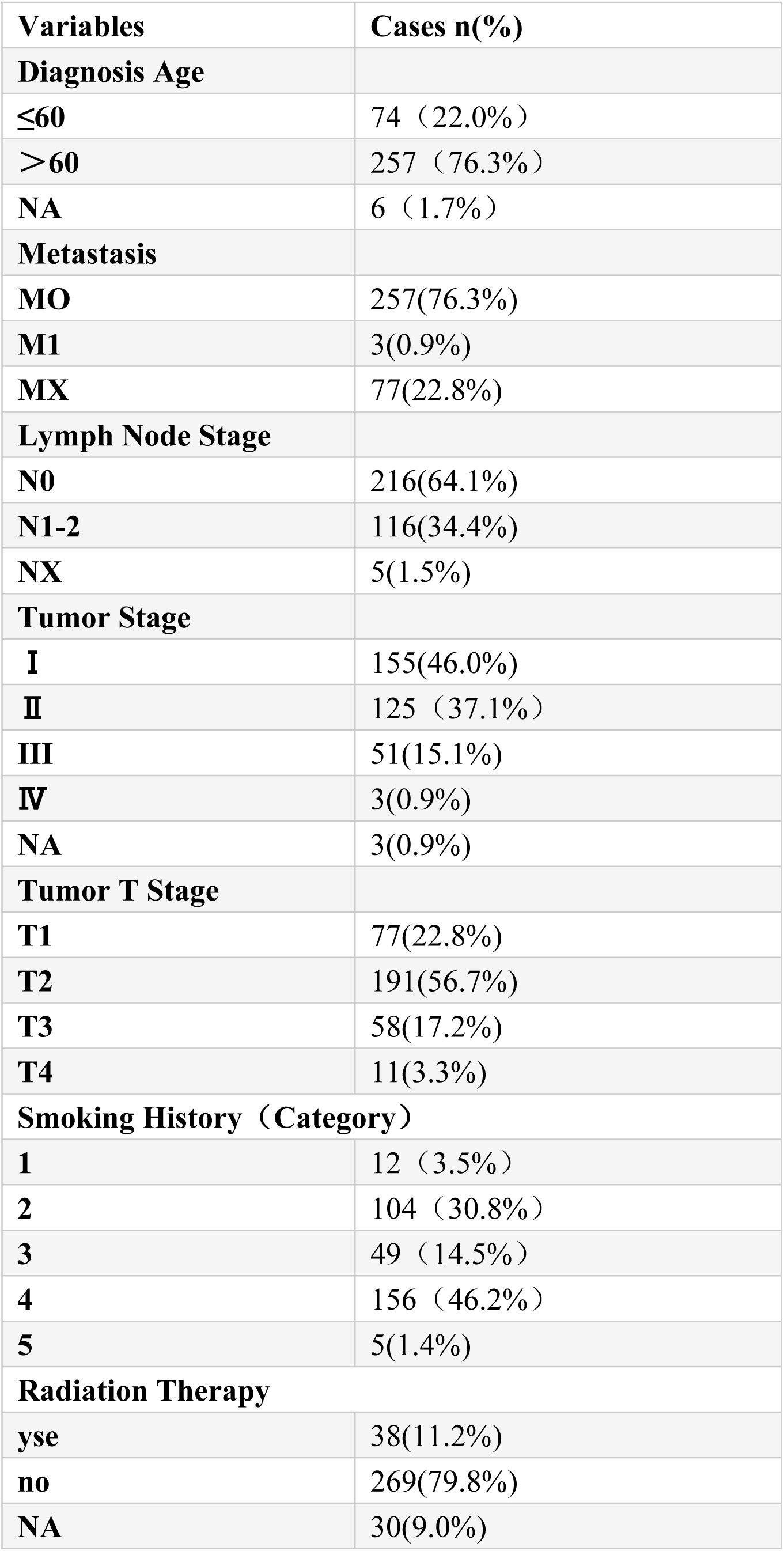
Clinical characteristics of LUSC

| **Variables** | **Cases n(%)** |
| --- | --- |
| **Diagnosis Age** |  |
| ≤60 | 74 (22.0%) |
| >60 | 257 (76.3%) |
| NA | 6 (1.7%) |
| **Metastasis** |  |
| MO | 257(76.3%) |
| M1 | 3(0.9%) |
| MX | 77(22.8%) |
| **Lymph Node Stage** |  |
| N0 | 216(64.1%) |
| N1-2 | 116(34.4%) |
| NX | 5(1.5%) |
| **Tumor Stage** |  |
| I | 155(46.0%) |
| II | 125 (37.1%) |
| III | 51(15.1%) |
| IV | 3(0.9%) |
| NA | 3(0.9%) |
| **Tumor T Stage** |  |
| T1 | 77(22.8%) |
| T2 | 191(56.7%) |
| T3 | 58(17.2%) |
| T4 | 11(3.3%) |
| **Smoking History (Category)** |  |
| 1 | 12 (3.5%) |
| 2 | 104 (30.8%) |
| 3 | 49 (14.5%) |
| 4 | 156 (46.2%) |
| 5 | 5(1.4%) |
| **Radiation Therapy** |  |
| yse | 38(11.2%) |
| no | 269(79.8%) |
| NA | 30(9.0%) |

**Table 2.**
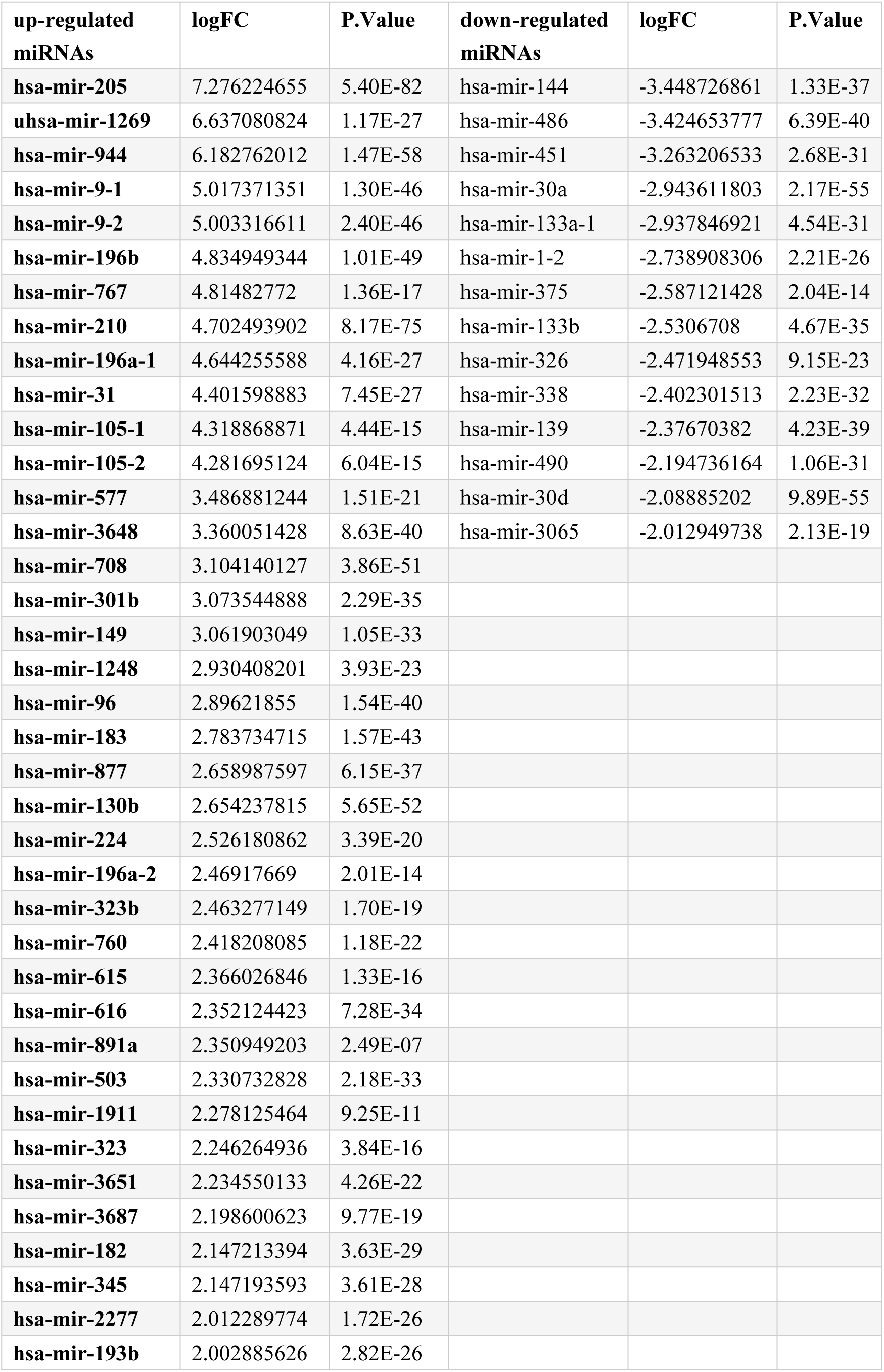
Differentially expressed miRNAs

### Dataset screening and verification of GEO database

A total of 4 eligible data sets were selected from the GEO database, including GSE16025, GSE19945, GSE51853 and GSE74190 (S3 Table). Due to the relatively small sample size of the dataset in the GEO database, our cut-off criteria is set at P<0.05 and |logFC|≥1. The differentially expressed miRNAs obtained from the four data sets were taken together (Fig. 2), and a total of eight differentially expressed miRNAs shared by four data sets were obtained. Then, the differentially expressed miRNAs obtained from the TCGA database were matched to and obtain four up-regulated miRNAs, hsa-mir-205, hsa-mir-210, hsa-mir-182, hsa-mir-224, and one down-regulated miRNA, hsa-mir-451.

**Fig 2.**
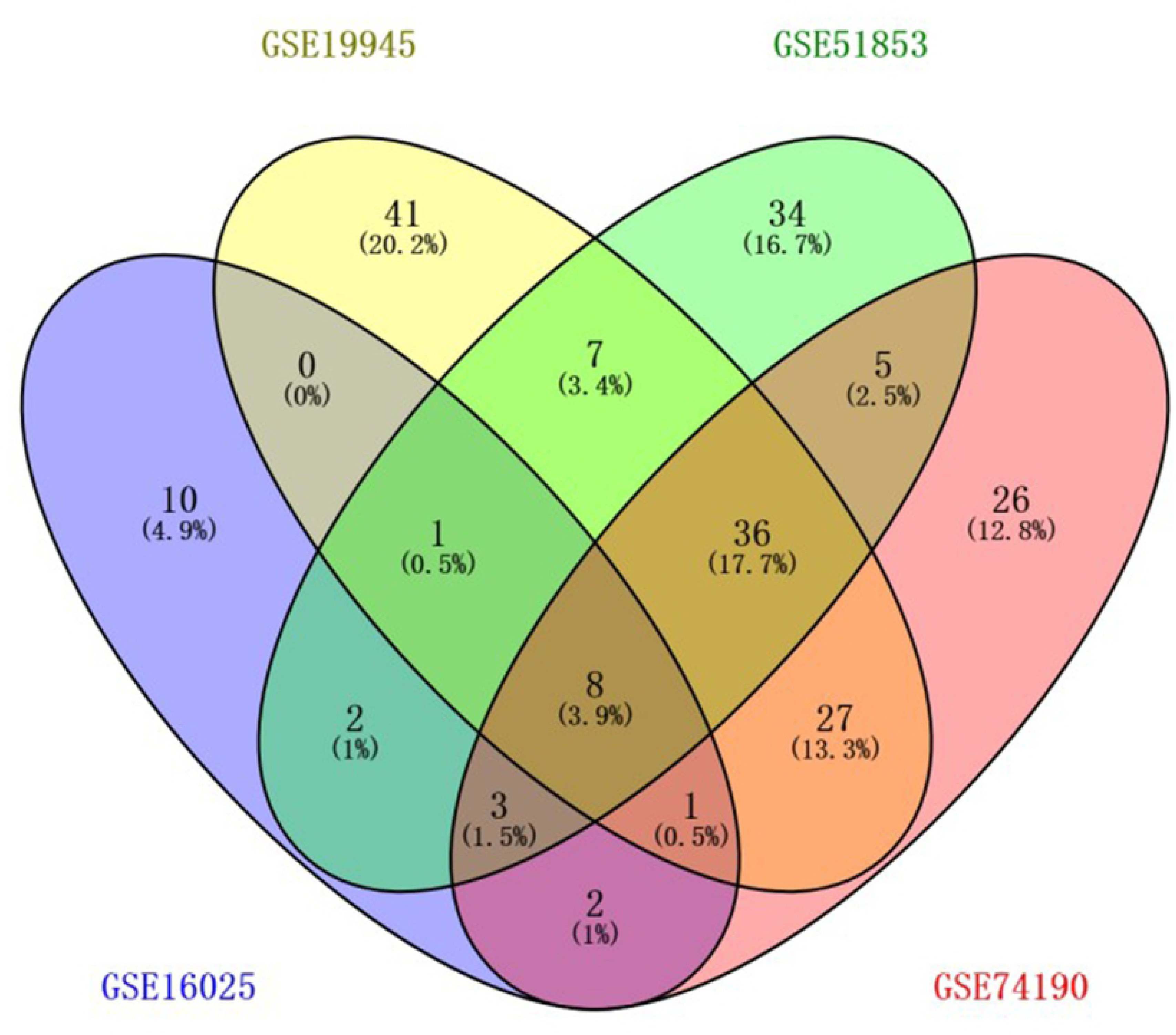
Venn diagram: differentially expressed miRNAs in 4 data sets.

### Prognostic value of differentially expressed miRNAs in LUSC

To identify the differentially expressed miRNAs which would be potentially associated with overall survival of LUSC patients, we evaluated the association between miRNAs expression and patients’ survival using Kaplan-Meier curve and Log-rank test. The results show that hsa-mir-205(P=0.367), hsa-mir-182(P=0.118), hsa-mir-224(P=0.555), hsa-mir-451(P=0.365) have no connection with OS. The hsa-mir-210(P=0.007) was negatively correlated with OS (Fig. 3). In the analysis of single factor clinical factors, Metastasis(P<0.001), T stage(P=0.0.012), and clinical stage(P=0.049) were also significantly associated with OS. In multivariate cox regression analysis, hsa-mir-210 and T staging were still significantly associated with OS (Table3). The overall survival of the hsa-mir-210 low expression group was significantly prolonged in both the early and late (T stage) LUSC patients. The hsa-mir-210 was showed to be an independent prognostic factor in LUSC. The association between hsa-mir-210 and clinical features was evaluated in LUSC patients, and was significantly associated with age (P=0.009), and the others were no significant difference with hsa-mir-210.

**Fig 3.**
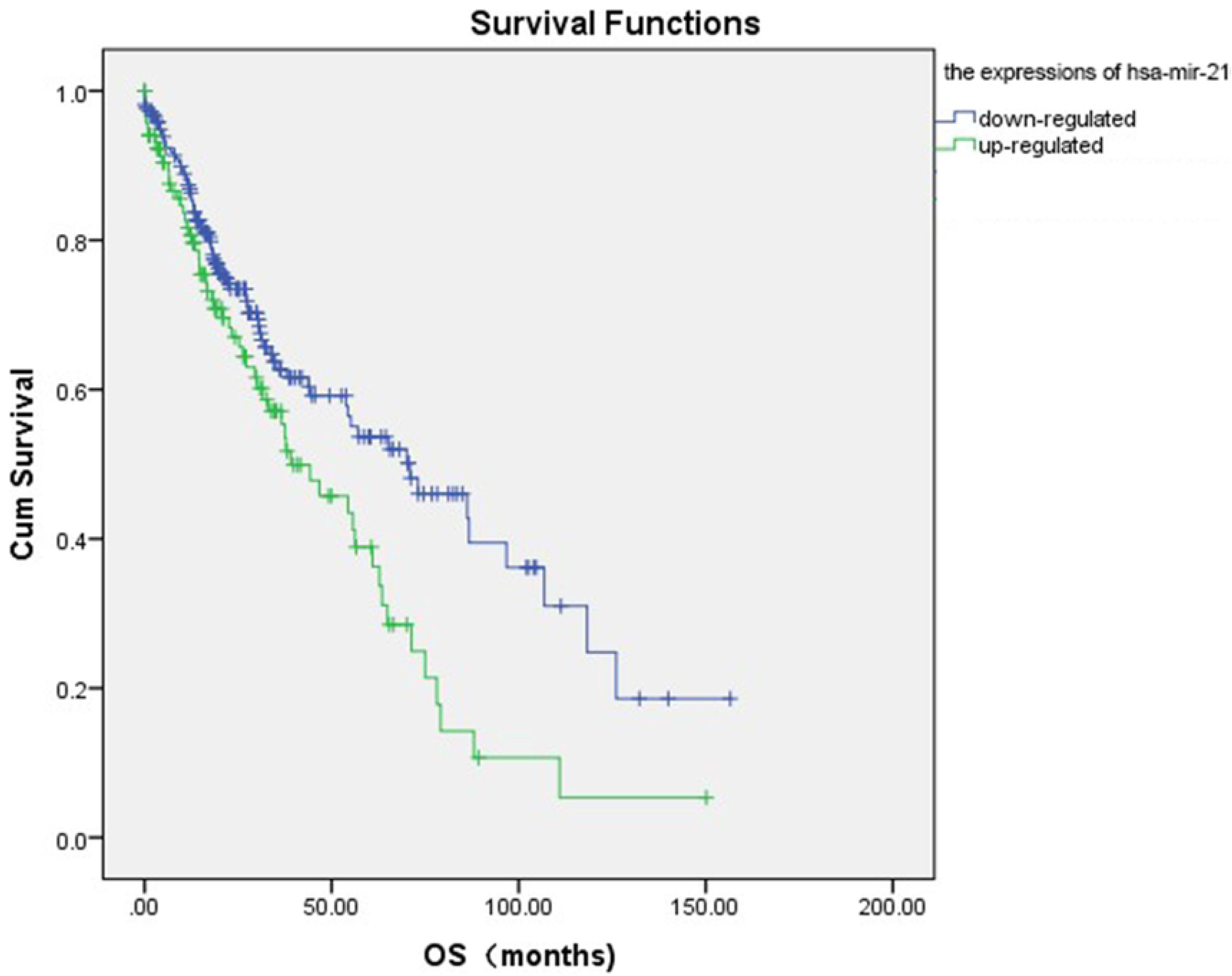
The survival functions of LUSC. The hsa-mir-210 was associated with overall survival in LUSC patients by using Kaplan Meier curve and Log-rank test. The patients were stratified into high level group and low level group according to average value.

**Fig 4.**
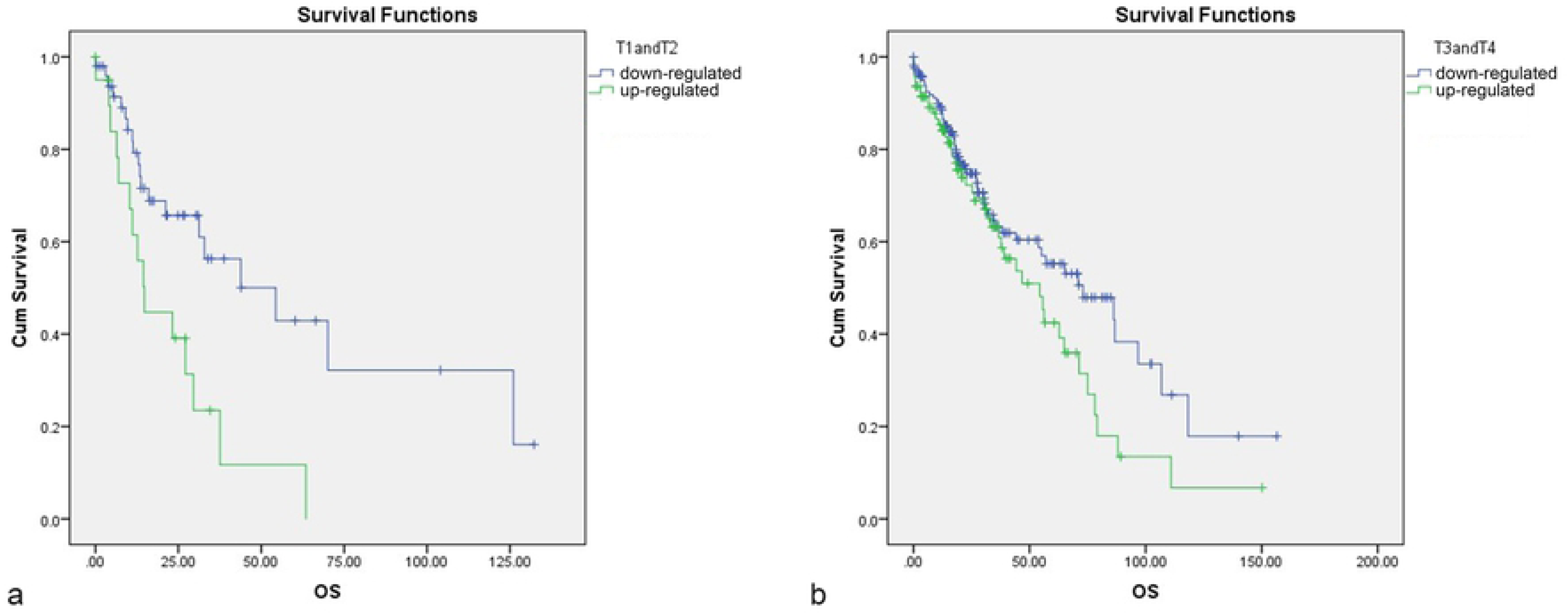
Relationship between miRNA expression and OS of different T stages: (a) early stage (T1-T2); (b) late stage (T3-T4).

**Table 3.** Univariate and multivariate Cox regression analysis in LUSC patients.

|  | Univariate analysis |  | Multivariate analysis |  |
| --- | --- | --- | --- | --- |
|  | HR (95%CI) | P.Value | HR(95%CI) | P.Value |
| Age(>60vs≤60) | 1.035(0.674-1.589) | 0.876 |  |  |
| Smoking history category(>3 vs ≤3) | 0.742(0.527-1.045) | 0.088 |  |  |
| Sex(male vs female) | 1.027(0.689-1.530) | 0.896 |  |  |
| Metastasis(M1 vs M0) | 7.912(2.476-25.288) | <0.001 |  |  |
| T stage(T3+T4 vs T1+T2) | 1.638(1.113-2.410) | 0.012 | 1.657(1.064-2.582) | 0.026 |
| Lymph node stage(N1-2 vs N0) | 1.068(0.754-1.511) | 0.713 |  |  |
| Radiation therapy(yes vs no) | 1.505(0.941-2.406) | 0.088 |  |  |
| Clinical stage(III+IV vs I+II) | 1.506(1.002-2.263) | 0.049 |  |  |
| hsa-mir-210(up-regulated vs down-regulated) | 1.529(1.134-2.233) | 0.007 | 1.725(1.216-2.448) | 0.002 |

### Target gene prediction and functional enrichment analysis

The target genes of hsa-mir-210 was predicted using TargetScan, miRDB, starBase, and miRanda online analysis tools. A total of 520 genes appearing in two or more databases were identified. (Fig. 5). The target genes are then functionally enriched to stage their biological functions. The KEGG pathways were significantly enriched in HIF-1 signaling pathway, cAMP signaling pathway, VEGF signaling pathway, MAPK signaling pathway, choline metabolism and Focal adhesion. The GO pathway were significantly enriched in negative regulation of cell differentiation, positive regulation of cell proliferation and positive regulation of autophagy, etc. (Fig. 6). The HIF-1 signaling pathway, MAPK signaling pathway and VEGF signaling pathway are interrelated and play a role in tumor growth and metastasis(Fig.7).

**Fig 5.**
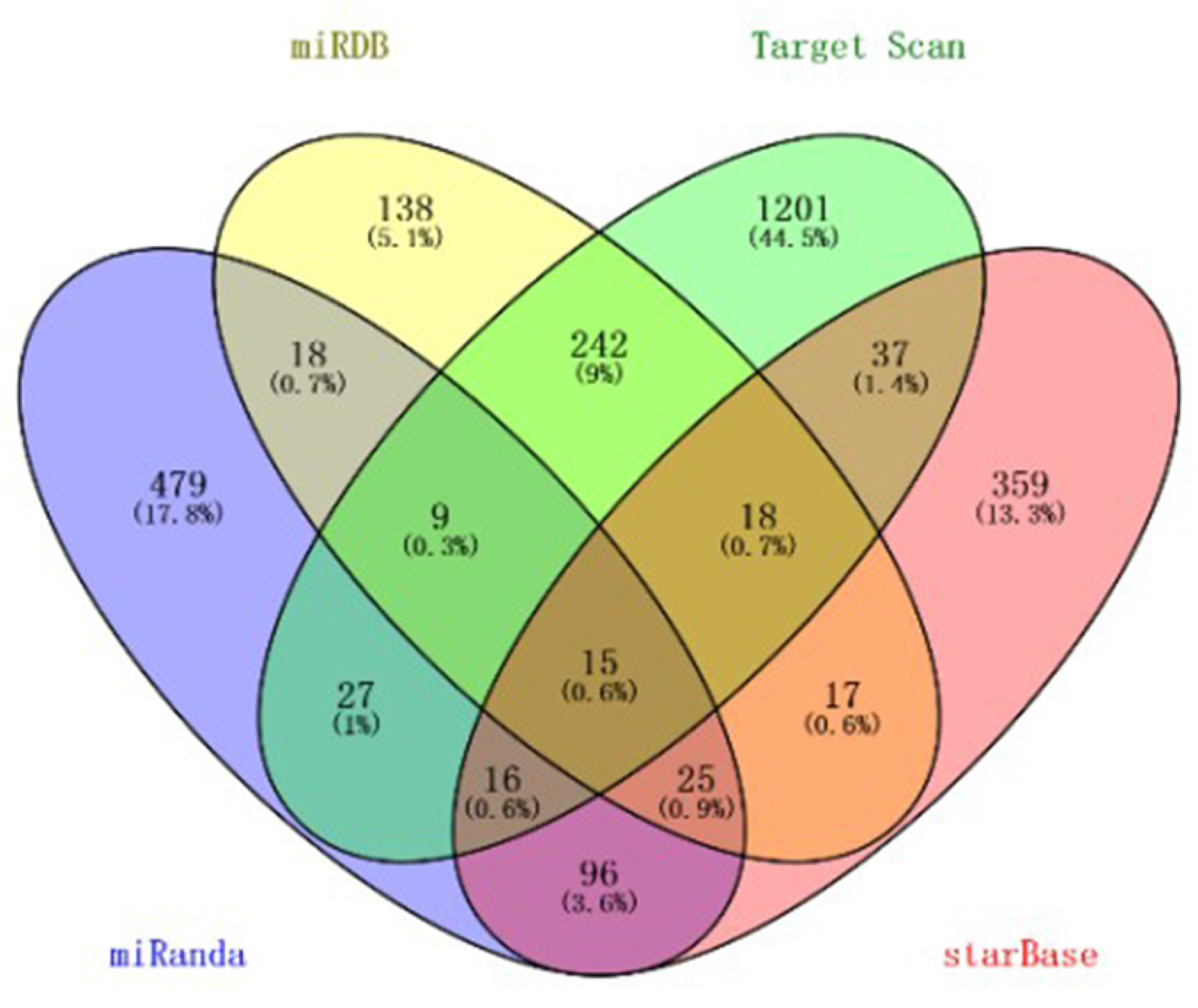
Venn diagram of target gene prediction.

**Fig 6.**
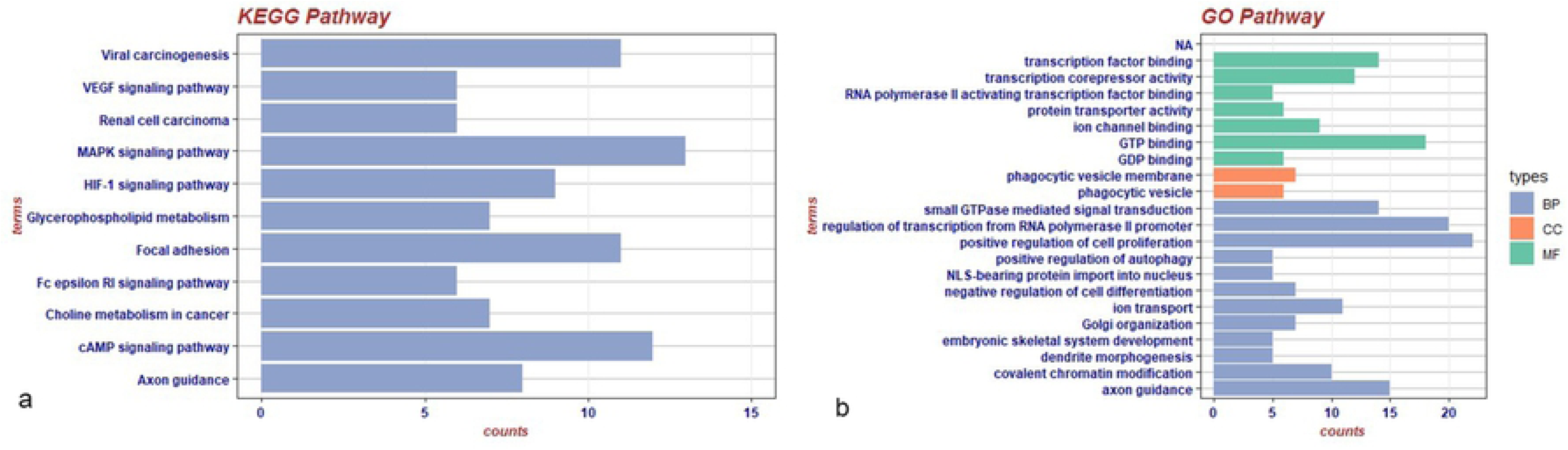
Functional enrichment diagram: (a) The enrichment results of KEGG Pathway;(b)The enrichment results of GO Pathway.

**Fig 7.**
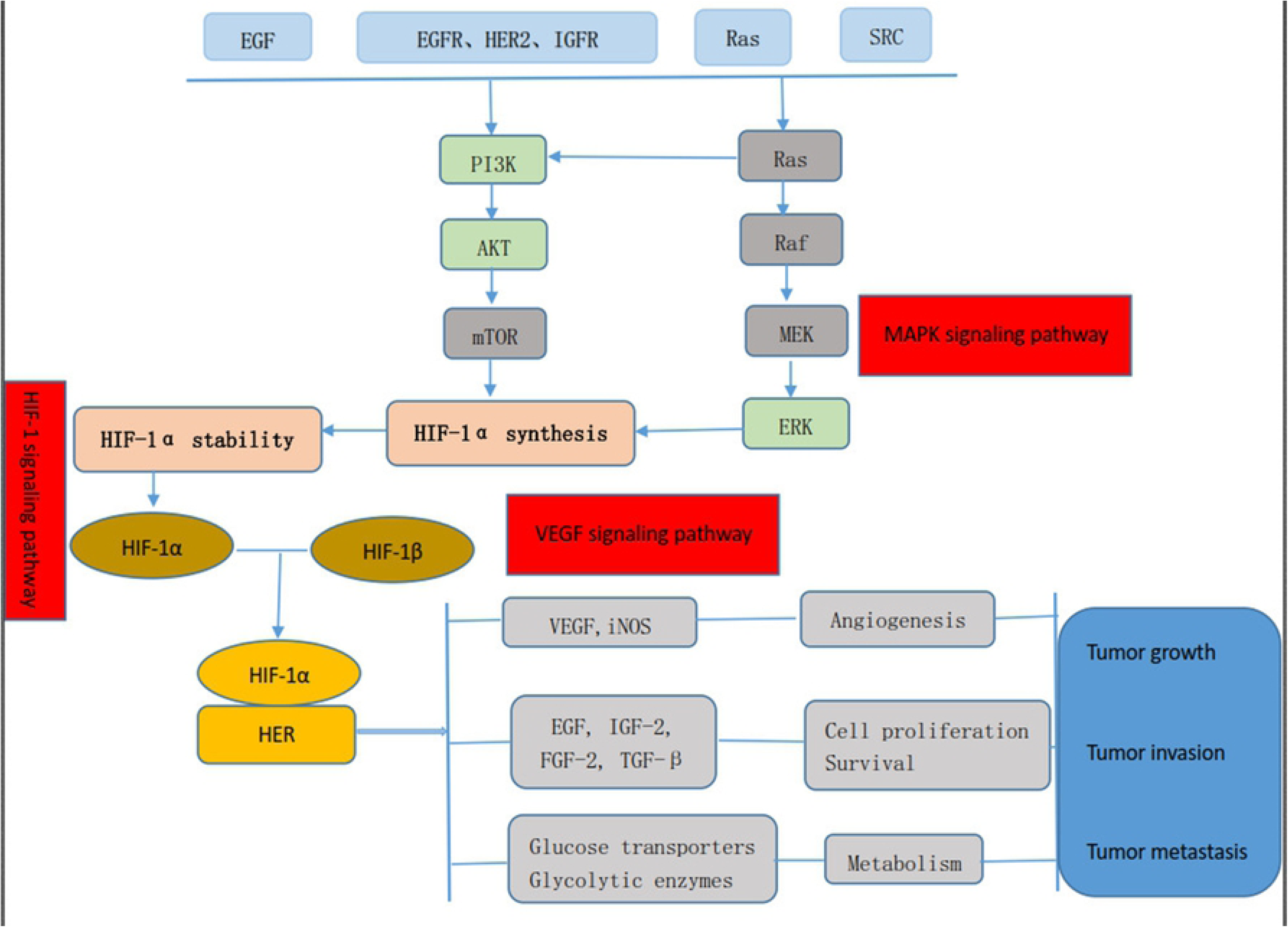
Pathway interaction diagram.

## Discussion

By 2015, 1.59 million people worldwide had died of lung cancer, of whom 30-40% had been diagnosed with NSCLC (squamous cell carcinoma). Lung cancer is also the leading cause of cancer death in China, and LUSC accounts for a high proportion. Because most LUSC patients are diagnosed at advanced stage, with high surgical risk and poor cardiopulmonary function, it has been considered as a refractory solid tumor [12-14]. If some markers can be found for early diagnosis, the diagnosis rate of patients will be greatly improved, and early intervention will improve the overall survival time of patients. In this study, we found five miRNAs (hsa-mir-205, hsa-mir-210,hsa-mir-182,hsa-mir-224,hsa-mir-451) that were significantly differentially expressed in tumor tissues and normal tissues through screening and validation, and one(hsa-mir-210) of them was significantly correlated with patient survival. After multivariate analysis, it was still significantly correlated with patient prognosis and was expected to be a new marker for predicting survival.

In the past decade, miRNAs have become biomarkers for diagnosis, prognosis and prediction of treatment response, both from tumor specimens and in biological fluids [15]. In terms of lung cancer, as early as in 2004 has made the role of miRNAs in lung cancer, there is evidence that more than half of microRNAs genes associated with cancer genome area or fragile sites, and the lack of a few miRNAs in lung cancer cell lines and the low level of expression of chronic lymphocytic leukemia [15, 16]. Subsequently, numerous studies have been published on the role of miRNAs in lung cancer. For example, the down-regulated expression of hsa-mir-199 in lung cancer is closely related to staging, distant metastasis and poor prognosis, and may inhibit the malignant progression of lung cancer by interacting with RGS 17[17]; hsa-mir-30e plays an inhibitory role in NSCLC, and may inhibit cell proliferation and invasion by directly targeting SOX 9[18]; The hsa-mir-451 regulates survival of NSCLC cells partially through the downregulation of RAB14 and targeting with the hsa-mir--451/RAB14 interaction might serve as a novel therapeutic application to treat NSCLC patients[19]. In addition, hsa-mir-373[20], hsa-mir-17-92 cluster [21, 22], hsa-mir-21[23], hsa-mir-126[24], hsa-mir-145[25], and hsa-mir-340[26] were also found to be closely related to the occurrence, progression and survival of lung cancer.

In this study, through the difference analysis of the TCGA database and the verification of the GEO database, a total of four miRNAs with significantly high expression (relative to normal tissue) were obtained, including hsa-mir-205, hsa-mir-210, hsa-mir-182, hsa-mir-224, and a low expression (relative to normal tissue) of hsa-mir-451. Moreover, the expression of hsa-mir-210 was also significantly correlated with patient survival [27]. Huang W et al. reported that the expression state of hsa-mir-205 could distinguish LUSC from LUAD and SCLC, and the diagnostic accuracy was relatively high. Jiang M et al. also confirmed the value of hsa-mir-205 in the diagnosis of NSCLC [28]. Zhu YJ et al. reported that hsa-mir-182 by inhibiting RASA1 expression to restrain cancer cell proliferation of lung squamous carcinoma[29], another study also showed that hsa-mir-182, hsa-mir-210 and other three miRNAs can be used as a new diagnostic markers for LUSC[30], and in stageII LUSC patients, high expression hsa-mir-182 is an independent positive prognostic factors[31]. There are relatively few reports on hsa-mir-224, but it could be used together with other miRNAs as an evaluation indicator for the palliative effect of advanced LUSC [32]. Studies of hsa-mir-451 have also been reported in NSCLC [33]. Taken together, 5 differentially expressed miRNAs provide a new direction for the diagnosis of LUSC, or may be used as diagnostic markers to improve the clinical diagnosis rate.

Subsequently, we analyzed the relationship between five differentially expressed miRNAs and OS, and found that the expression level of hsa-mir-210 was significantly correlated with OS [34, 35], however, the relationship between the expression of hsa-mir-210 and the survival of LUSC has rarely been reported. In this study, we obtained through KM survival analysis that the overall survival of patients with low expression of hsa-mir-210 was significantly prolonged in the higher expression group, and the difference between the two groups was statistically significant (P=0. 007). In the subsequent univariate and multivariate analyses, the P values were 0.007 (HR= 1.529, 95%CI: 1.134-2.233) and 0.002(HR=1.725, 95%CI: 1.216-2.448), which were still statistically significant. At the same time, T staging of the tumor was also significantly associated with survival in multivariate cox analysis. We then analyzed the relationship between the expression of hsa-mir-210 and survival in different T staging subgroups, and found that in the early (T1, T2) and late (T3, T4) T staging subgroups, the results showed that the overall survival of patients in the low expression group of hsa-mir-210 was significantly better than that in the high expression group. Based on the above analysis results, it can be seen that hsa-mir-210 is an independent prognostic risk factor affecting the survival of patients with LUSC and can be used as a potential prognostic factor.

In order to further investigate hsa-mir-210, we predicted its target gene and then analyzed the pathway and related functions. In the analysis, hsa-mir-210 was closely related to positive and negative regulation of cell proliferation, DNA transcription, VEGF signaling pathway, MAPK signaling pathway, hif-1 signaling pathway and choline metabolism pathway. Moreno Roig et al. reported that the expression of HIF-1 in NSCLC has the effect of increasing radio sensitivity [36];The MAPK signaling pathway has also been reported in cell proliferation, differentiation, migration, aging and apoptosis[37]; Glunde K et al. also made a detailed report on the role of choline metabolism in the diagnosis of tumor molecules[38]. Therefore, we need to further study the molecular mechanisms of these pathways to provide new clinical interventions for LUSC to improve patient survival. And the pathway interaction also reveals the role of hsa-mir-210 in the growth, invasion, and metastasis of LUSC

To sum up, we analyzed 5 potential miRNAs that may be signaling molecules for the diagnosis of LUSC, and identified hsa-mir-210 that may be a potential prognostic factor for LUSC. Further studies need to validate our findings in large samples, and further functional studies also need to explore the molecular mechanisms of these miRNAs in the progression of LUSC.

## Materials and Method

### Data download and processing

In this study, raw miRNAs expression data from patients with LUSC came with http://firebrowse.org/LUSC miRseq (illuminahiseq_mirnaseq_gene_expression.MD5), a total of 387 samples of miRNAs expression information, including 342 tumor samples, 45 normal tissue samples. The clinical information of the sample was downloaded from the cBioPortal website (lung squamous cell carcinoma, TCGA Provisional, clinical data) and matched with the miRNAs data samples. A total of 337 samples with complete clinical information and miRNAs expression information were obtained. The data were processed by RStudio (Version 1.1.463, limma package). A total of 1047 miRNAs were involved in the analysis. Under the condition of P<0.05, the expression was up-regulated by logFC≥2, and the expression of logFC≤-2 was down-regulated.

### Screening and Analysis of Validation Datasets in GEO Database

Search the relevant database on the GEO website (https://www.ncbi.nlm.nih.gov/geo/) and search for the keywords (microRNA OR “miR” OR “miRNAs”) AND (Lung OR pulmonary) AND (Cancer OR tumor OR neoplasm OR malignancy OR carcinoma OR SCC OR NSCLC), select “series” for the data type, “Homo sapiens” for the species. For data sets related to NSCLC, the inclusion criteria for the data set are: 1) the sample is diagnosed as NSCLC and the pathological type is squamous cell carcinoma; 2) the cancerous tissue specimen and the lung cancer tissue specimen (healthy human lung tissue or cancer) Next to normal lung tissue); 3) more than 5 tumor samples and normal samples; 4) is the expression data spectrum for sequencing the miRNAs of the sample; 5) The R online tool can be used. Then the GEO analysis tool R was used to obtain the difference analysis data of the four data sets respectively. Under the condition of P<0.05, the |logFC| ≥1 was used as the screening standard. Based on the TCGA database difference analysis data and the validation data of the GEO dataset, four high-expression miRNAs were obtained, namely: hsa-mir-205, hsa-mir-210, hsa-mir-182, hsa-mir-224; Low expression of miRNA, hsa-mir-451.

### Analysis of relationship between differentially expressed miRNAs and patient prognosis

Differentially expressed miRNAs were divided into high expression group and low expression group according to the average level of expression, using Kaplan-Meier (K-M) survival analysis and Log-Rank test of SPSS (IBM SPSS Statistic 23). The effect of clinical factors on miRNAs expression was tested by independent sample t test. The univariate and multivariate Cox regression analysis were used to test the effect of the miRNAs and clinical factors on OS.

### Target gene prediction and functional enrichment analysis of prognostic miRNA

The target genes are predicted at miRanda (http://www.microrna.org/microrna/home.do), miRDB (http://mirdb.org/miRDB/), TargetScan (http://www.targetscan.org/), starBase (http://starbase.sysu.edu.cn/) online tools. The prediction results are displayed using Wayne Chart (VENNY2.1). Then, genes that overlap in two databases or above were analyzed by The Database for Annotation, Visualization and Integrated Discovery (DAVID) bioinformatics tool (https://david.ncifcrf.gov/). DAVID is a bioinformatics database that integrates biological data and analysis tools to provide systematic and comprehensive bioinformatics annotations for large-scale gene or protein lists (hundreds or thousands of gene ID or protein ID lists) to help users extract bioinformatics from them. Kyoto Encyclopedia of Genes and Genomes (KEGG) pathway and Gene Ontology (GO) enrichment analyses were then performed for the target genes. The P-value<0.05 and gene count≥5 were set as the cut-off criteria.

## Acknowledgments

First of all, I would like to express my deepest gratitude to my graduate tutor, Professor Xia Shu. From the design idea of this article to the final draft, Professor Xia gave valuable Suggestions in every link, and I would like to thank Professor Xia for his patient guidance. Secondly, I would like to thank teacher Chen Yu for her suggestions on the structure of my article. Thanks for the convenient platform provided by Tongji Hospital, Tongji Medical College, Huazhong University of Science and Technology. Last but not least, I would like to thank my senior sister and junior sister for their support in my work. In particular, thanks to Shuai-wen Huang for polishing and typesetting my article.

## Supporting information

**S1 Table. Original data table of sample miRNAs expression.**

**S2 Table. Raw data table of clinical information of the sample**

**S3 Table. The information of GEO data**

